# Summary of muscle parameters for Hill-based muscle modeling in the human lower limb

**DOI:** 10.1101/090944

**Authors:** Ross H. Miller

## Abstract

A summary is presented of five mechanical parameters from human lower limb skeletal muscles critical for Hill-based muscle modeling: the optimal fiber length, the fiber pennation angle, the physiological cross-sectional area (PCSA), the unloaded tendon length, and the fast-twitch fiber fraction. The data presented are drawn from a total of 29 publications including human cadaver studies, *in vivo* imaging studies of live humans, musculoskeletal modeling studies, and combinations of these methods. Where possible, parameter values were adjusted from the referenced data to present them with consistent definitions (normalization of measured fiber lengths to optimal sarcomere length, and calculation of PCSA as the ratio of fiber volume to fiber length). It is seen that within a specific muscle, optimal fiber lengths are fairly consistent between studies, pennation angles and PCSAs vary widely between studies, and data for unloaded tendon length are comparatively sparse. Few studies have reported fiber type fractions for a large number of muscles. Guidelines for implementing these parameter values in muscle modeling and musculoskeletal modeling are suggested.

**Update History:**

1. December 2, 2016: original submission
2. December 3, 2016: discussion on maximum isometric force and specific tension added
3. December 6, 2016: minor edits for typos, clarity, and missing references
4. December 7, 2016: Tirrell et al. (2012) study added

## Introduction

The Hill muscle model (Fig. 1) consisting principally of a contractile component (CC) in series with an elastic component (SEC) was first posed conceptually by Hill (1938), with the earliest implementation in computer simulations of human movement by Hatze (1976). The Hill model today is used widely as an actuator in musculoskeletal modeling software such as SIMM, OpenSim, AnyBody, LifeMOD, and various custom models (e.g. Koelewijn & van den Bogert, 2016). Using the Hill model in practice requires specification of numerous parameter values such as the maximum isometric force, the optimal CC length, and the unloaded SEC length. When simulating the actions of a specific muscle or a whole-body motion actuated by many muscles, it is important for these parameter values to be defined on an appropriate muscle-specific basis (Caldwell & Chapman, 1989; Pandy, 1990; Caldwell, 1995; Scovil & Ronsky, 2006; Redl et al., 2007; De Groote et al., 2010; Carbone et al., 2016). Inaccurate or unrealistic parameter values can affect model-based predictions of, for example, knee joint loading during walking (Navacchia et al., 2016) and height achievement in vertical jumping (Domire & Challis, 2010). The process of assigning many parameters for many muscle models can be daunting, especially for new users who may be unfamiliar with the need, available resources, and methods for doing so. Prior summaries of mechanical parameters for individual human muscles exist, but are available only in textbooks (Yamaguchi et al., 1990; van der Helm & Yamaguchi, 2000). The most recent was published 16 years ago, with several landmark studies occurring in the intervening time (e.g. Klein Horsman et al., 2007; Ward et al., 2009), and no summary to date has included guidelines for implementation of the parameter values in muscle modeling.

**Figure 1.**
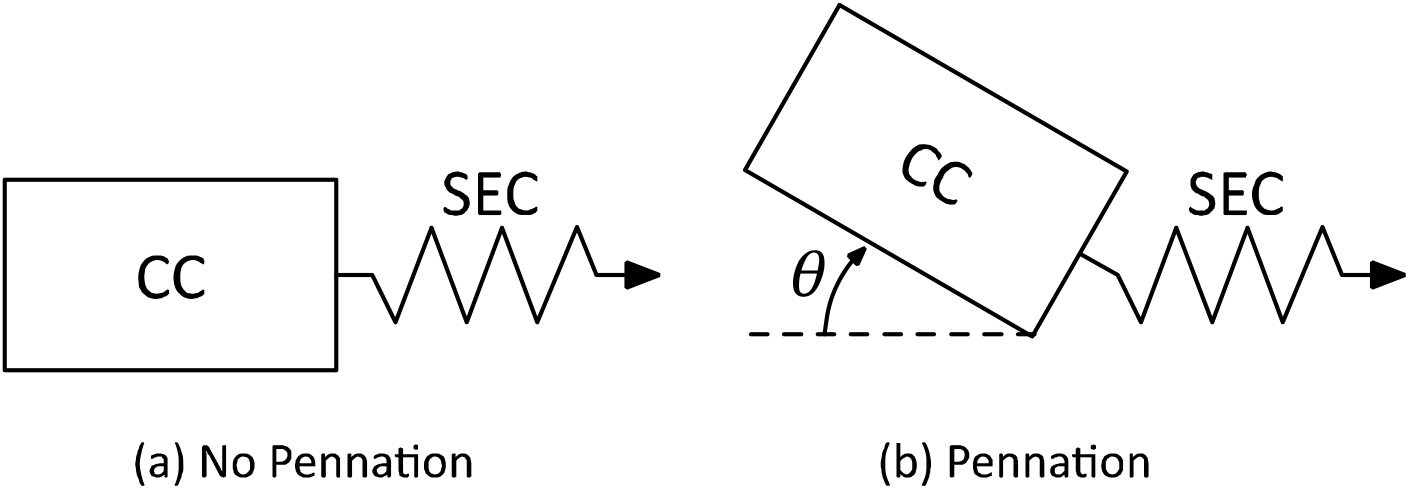
The two-component Hill muscle model with a contractile component (CC) in series with an elastic component (SEC). Model (a) has no CC pennation; Model (b) includes CC pennation with angle *θ*. The CC produces active force in response to an input excitation signal and its current length and velocity. The CC expresses this force across the SEC, which responds by changing length, potentially changing the length and velocity of the CC depending on the whole-muscle kinematics. The pennation angle *θ* changes to keep the CC volume constant.

Therefore, the purpose of this study was to summarize the available data on mechanical parameters of individual human lower limb muscles for Hill-based muscle modeling, and to suggest guidelines for their use and implementation in musculoskeletal modeling and computer simulation of human movement.

## 2. Methods

### 2.1 Studies and Parameters Included

The studies included were based on review of previous similar summaries (Yamaguchi et al., 1990; van der Helm & Yamaguchi, 2000), on recent related studies in the intervening years of 2000-2016 (e.g. Klein Horsman et al., 2007; Ward et al., 2009), and on manual review of studies that included at least one of the muscles from the comprehensive list of Klein Horsman et al. (2007). The PubMed and Google Scholar search engines were used.

The parameters considered were the optimal fiber length (*L*_*o*_), the fiber pennation angle (*θ*), the physiological cross-sectional area (PCSA), the unloaded tendon length (*L*_*u*_), and the fraction of “fast-twitch” fibers *FT.* The former four parameters were selected due to their frequency of appearance in the reviewed literature and the high sensitivity of Hill model output to their values (e.g. Scovil & Ronsky, 2006). Data on *FT* were included due to their importance in some models of muscle energy expenditure (e.g. Umberger et al., 2003).

A comment on terminology is warranted before proceeding. When using the terms “fiber” and “tendon” in the context of measurements from real muscles and Hill-based muscle modeling, the reader should keep in mind that the Hill model is *phenomenological* in nature: its purpose is to accurately simulate muscle force production for a variety of input excitation conditions and *whole-muscle* (origin-to-insertion) kinematic states. The CC and SEC in the Hill model capture many of the functions typically attributed to fibers and tendons respectively in real muscle, but the CC and SEC do not have direct anatomical analogues in real muscle. Hill himself cautioned against an anatomical interpretation of the Hill model’s components:

> “For simplicity in description the [SEC] will be referred to as ‘tendon’, but no assumption is implied that other undamped series elastic elements do not exist within the fibers themselves; the evidence of its properties is derived from mechanical experiments with active muscle, not from histological observation.” (Hill, 1950)

This caution has implications in how the parameters presented here are used in practice in muscle modeling. Guidelines for their implementation are a focus of the Discussion section.

### 2.2 Optimal Fiber/CC Length

The optimal fiber/CC length *L_o_* is defined as the length at which the fibers/CC produce the maximum isometric force *F_o_* when maximally activated at zero velocity. Methods of determining *L_o_* from measurements have varied in the literature. Some studies have determined fiber length from manual ruler-based measurements. In these cases, it is often the fascicle length (not the fiber length) that is measured, with the assumption that the fibers run the full length of the fascicles. This distinction can be largely semantic depending on the use of the data, but the assumption of equal fiber and fascicle lengths is controversial particularly for long fascicles (Trotter, 1990). Another common method of determining *L_o_* has been to report fiber lengths that have been “normalized” using sarcomere length measurements:

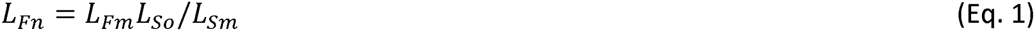

where *L_Fn_* is the normalized fiber length, *L_Fm_* is the originally measured fiber length, *L_So_* is the optimal sarcomere length in the context of the sliding filament hypothesis and the force-length relationship (Ramsey & Street, 1940; Gordon et al., 1966), and *L_Sm_* is the measured sarcomere length. The purpose of normalization is to obtain an estimate of optimal fiber length *L_o_*, i.e. *L_Fn_* ≈ *L_o_*. Such an estimate is necessary to produce the most accurate calculations of the PCSA, where *L_o_* appears in the denominator of the calculation (see §2.4). Fibers are not necessary at their optimal lengths during measurement, fiber operating lengths can vary by ±50% or more of *L_o_*, and measurements of fiber length at an unknown position on the force-length curve could thus introduce large errors in calculations of PCSA. The obvious challenge in normalization is that it requires measurements of sarcomere length. The optimal sarcomere length for human muscle appears to be about 2.6-2.8 *μ*m (Walker & Schroedt, 1974; Lieber et al., 1994). Unless otherwise noted, all fiber lengths that were normalized to an optimal sarcomere length other than 2.7 *μ*m in the referenced data were re-normalized to an optimal length of 2.7 *μ*m for presentation of the data here.

Another method for determining *L_o_* has been to use muscle modeling and optimization to adjust the value of *L_o_* (and various other model parameters) to track measurements of human joint torque production as well as possible. For example, Hasson and Caldwell (2012) optimized *L_o_* for gastrocnemius, soleus, and tibialis anterior to track measured maximum isometric torque-angle data from the ankle joint on a dynamometer. Relatedly, as noted in the previous section (§2.1), direct assignment of CC parameter values like *L_o_* from measurements on muscle fibers is against the spirit and purpose of the Hill model and should be avoided.

### 2.3 Fiber Pen nation Angle

Fiber pennation angle *θ* is typically defined as the orientation of the long axis of the muscle fibers or fascicles, relative to the tendon. This definition has been used fairly consistently in the literature. Cadaver studies by necessity measure *θ* when muscles are inactive. *In vivo* studies have sometimes measured *θ* at rest or during active contractions, and at various joint angles. These factors should be considered when interpreting *in vivo 9* data because length and activation level will affect the current *θ*.

Most studies have measured *θ* from a single plane and report a representative or average result for the muscle in question. This definition of *θ* is likely not reflective of the full fiber geometry in a three-dimensional muscle, but it is consistent with how *θ* is usually included in Hill-based models, where constant muscle thickness and volume is assumed. Infrequently, more detailed multi-planar definitions of *θ* have been used, and occasionally the distribution of pennation angles within the fibers of the muscle have been assessed (e.g. Scott et al., 1993).

### 2.4 Physiological Cross-Sectional Area

PCSA has been defined inconsistently in the literature. Some studies use the definition of Alexander and Vernon (1975):

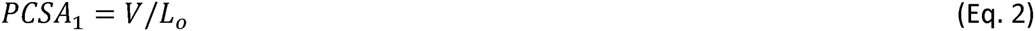

where *V* is the total volume of fibers within the muscle. This definition is advantageous because it has an intuitive geometrical interpretation (the cross-sectional area of contractile material perpendicular to the long axis of the muscle fibers) and because the product with the specific tension gives the maximum isometric force of the CC. An alternative definition was proposed by Sacks and Roy (1982):

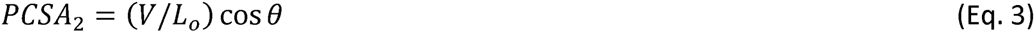

With this definition of PCSA, the product with specific tension gives the fraction of the CC maximum isometric force that can be expressed across the tendon at pennation angle *θ*. Equations 2 and 3 obviously produce identical results when *θ* is zero. Equation 2 was used for reporting PCSA in this study because (i) CC maximum isometric force is an important parameter in Hill-based muscle models, (ii) *θ* varies with muscle length and force and these factors are not controlled consistently between studies, (iii) some implementations of the Hill model do not include fiber pennation, and (iv) some studies using Eq. 2 have not measured or reported *θ*, making accurate conversions impossible.

With both definitions of PCSA, note that the result will be inaccurate when a fiber length other than *L_o_* is used in the denominator. Thus, values of PCSA reported from calculations involving non-normalized fiber lengths should be interpreted with caution if they are to be used to calculate the CC maximum isometric force.

### 2.5 Unloaded Tendon/SEC Length

Measurement of *L_u_* is a complicated endeavor because the “tendon” of a muscle can be a difficult structure to identify and define with consistency between muscles. In some muscles the tendon may run the full length of the muscle. The portion of the tendon that muscle fibers directly attach to is typically referred to as the *internal tendon* and may be fully or partially aponeurotic. The remaining portion of the tendon with no direct fiber attachments is typically referred to as the *external tendon.* The external tendon is the closest analogue to the Hill model’s SEC, but some muscles have two external tendons, one distal and one proximal to the fibers. For this reason and others (see §2.1), direct assignment of *L_u_* from measurements on tendon are discouraged.

Cadaver studies can measure tendon length when the muscles have been excised from the body, assuring a truly unloaded condition. With *in vivo* imaging studies, it is difficult to ensure that the tendon is truly unloaded. Even in the absence of active force production, some passive force may be transmitted through the tendon.

### 2.6 Fiber Type Distribution

Methods of defining and determining muscle fiber type vary widely and are beyond the scope of this review. The parameter *FT* is used here with the acknowledgement that this binary classification scheme may be overly simplistic. Interested readers are referred to the summary by Scott et al. (2001). Readers should note that the great majority of data on muscle fiber types is expressed as the fractions of fiber numbers. Fiber types have less frequently been expressed as the fractions of cross-sectional area. This distinction is important because single “fast-twitch” fibers tend to be larger than single “slow-twitch” fibers. Flowever, data from Clarkson et al. (1980) and Parkkola et al. (1993) suggest this distinction might only affect *FT* fractions by up to 10%.

### 3. Results

#### 3.1 Cadaver Studies of Many Muscles

Five studies with cadaver data from numerous muscles of the lower limb were identified; in order of publication year: Alexander and Verson (1975) presented data from the knee and ankle muscles of one cadaver; Wickiewicz et al. (1983) presented data from the knee and ankle muscles of three cadavers; Friedrich and Brand (1990) presented data from the hip, knee, and ankle muscles of two cadavers; Klein Florsman et al. (2007) presented data from the hip, knee, and ankle muscles of one cadaver; Ward et al. (2009) presented means from the hip, knee, and ankle muscles of 21 cadavers. Alexander and Vernon (1975) and Friedrich and Brand (1990) presented raw measured fiber lengths. The other studies all presented normalized fiber lengths. Fiber lengths from Wickiewicz et al. (1983) were originally normalized to an optimal sarcomere length of 2.2 *μ*m and were renormalized to 2.7 *μ*m in the values presented here, with the PCSAs recalculated accordingly. Among these five studies, Friedrich and Brand (1990) is unique in the presentation of data from a male cadaver of relatively young (age 37 years), Klein Florsman et al. (2007) presents data from the most muscles and is the only study to present tendon length data, and Ward et al. (2009) has by far the largest sample size, with three times the sample size of the other four studies combined. In cadaver studies, muscle fiber volume for use in Eq. 2 has typically been determined by measuring muscle mass after stripping away tendon and connective tissue, and assuming a density of about 1.06 g/cm^3^ (e.g. Mendez & Keys, 1960).

A cadaver study by Weber (1846) is cited frequently in this literature, but the author has been unable to find an original copy of this study. Data from Weber (1846) are presented in Friedrich and Brand (1990), but the data are incomplete and the PCSAs have been scaled by an unknown constant.

#### 3.2 Cadaver Studies of a Small Number of Muscles

Three cadaver studies that presented data on a small number of muscles were identified. Scott et al. (1993) presented volume data for 14 muscles from a single cadaver, but data for fiber length and PCSA were only available for semimembranosus and vastus medialis. Raw measured fiber lengths were used in the PCSA calculations. Woodley and Mercer (2005) performed a detailed anatomical investigation of the hamstrings muscles in six cadavers, with PCSAs calculated using raw measured fiber lengths. The values for *L_u_* reported here are the sum of the proximal and distal external tendon lengths. Regev et al. (2011) presented data from 14 cadavers on psoas major, using measured sarcomere lengths for normalization of *L_o_*

#### 3.3 In Vivo Imaging Studies

Six studies used medical imaging modalities to estimate muscle parameters *in vivo* in living human subjects. Among these studies, Akima et al. (2001) presented data for the largest number of muscles (14) in a sample of 15 young male subjects, although the only parameter presented was PCSA. Fiber volumes were measured from cross-sectional MRI slices, and fiber lengths were estimated from muscle length assuming the same ratios of fiber-to-muscle length reported by Wickiewicz et al. (1983). This approach is common in MRI studies because (i) measurement of fiber/fascicle lengths is difficult with this imaging modality and (ii) technology for non-invasive measurement of sarcomere lengths has only recently been developed (Sanchez et al., 2015). The mean PCSAs reported by Akima et al. (2001) were adjusted here to account for the fact that Wickiewicz et al. (1983) calculated fiber lengths normalized to an optimal sarcomere length of 2.2 *μ*m rather than 2.7 *μ*m.

Maganaris et al. (2001) presented data for soleus and tibialis anterior in six young male subjects. Fiber volumes were measured from MRI and fascicle lengths were measured from ultrasound during maximum voluntary isometric contractions. Fukunaga et al. (1992) presented data on seven ankle muscles plus popliteus in 12 young adults, using similar methods to Akima et al. (2001). Reeves et al. (2004) presented data on vastus lateralis in 18 older adults using ultrasound to measure fiber volumes and fascicle lengths during maximum stimulated isometric contractions. Narici et al. (2003) presented data on medial gastrocnemius in 14 young males and 16 older males. Fiber volumes were measured from CT scans and fascicle lengths were measured at rest using ultrasound. O'Brien et al. (2010) reported data for the quadriceps in 20 young adult subjects. Fiber volume was measured using MRI and fascicle lengths were measured during maximum voluntary isometric contractions using ultrasound, at the knee joint angle that elicited the greatest extensor torque.

#### 3.4 Musculoskeletal Modeling Studies

If the eventual goal of obtaining muscle parameters is to use them in a musculoskeletal model, it can be informative to include the musculoskeletal model itself in the process of determining the parameter values. In a recent example, Arnold et al. (2010) used mean cadaver data from Ward et al. (2009) to update an earlier musculoskeletal model (Delp et al., 1990). Values for the unloaded tendon lengths were calculated by subtracting the Ward et al. (2009) fiber lengths from the origin-to-insertion muscle lengths of the model when its joints were in the same position as the joints of the cadavers during the Ward et al. (2009) measurements. The model’s active and passive torque-angle relationships were then compared to normative human data, with generally favorable results. The Arnold et al. (2010) parameters are presented in the spreadsheet combined with the Ward et al. (2009) parameters due to their close association. Values for PCSA from Arnold et al. (2010) that were not included in Ward et al. (2009) were calculated by dividing the maximum isometric force by a specific tension of 61 N/cm^2^, which was the value used by Arnold et al. (2010) to convert the Ward et al. (2009) PCSAs to forces.

A more complicated modeling approach was applied to the ankle joint by Hasson and Caldwell (2012). Maximum voluntary isometric contractions of plantarflexion and dorsiflexion were performed at a range of joint angles by 12 young adults and 12 older adults. A planar model of the ankle joint including soleus, gastrocnemius, and tibialis anterior was used to simulate these contractions for each subject, and the optimal CC length *L_o_* and unloaded SEC length *L_u_* (along with other parameters) were optimized to track the measured torque-angle data. In the results presented here, mean fiber volumes from MRI were divided by mean *L_o_* to determine the reported PCSA values. An advantage of this approach is that it is faithful to the spirit of the Hill model: parameters are defined such that joint-level output resulting from muscle forces is realistic. A disadvantage is that there can be a mismatch between the sources of the model’s output and the sources of the measurements. For example, plantarflexors other than the triceps surae complex and dorsiflexors other than tibialis anterior contributed to the measured torque, but were not modeled.

A large number of musculoskeletal models have based their muscle model parameter values partly or entirely on the influential lower limb model by Delp et al. (1990), which used similar methods to Arnold et al. (2010) and referenced many parameters from the young male cadaver of Friedrich and Brand (1990). To avoid redundancy in the data presented here, models with parameter values based on Delp et al. (1990) were excluded with the exception of the most recent iteration of the original model, the popular “Gait2392” model available in OpenSim software (Thelen & Anderson, 2006; Delp et al., 2007). Specific tension of 61 N/cm^2^ was again used to convert maximum isometric force to PCSA; Delp et al. (1990) used this value in the original derivation of the model’s parameters.

#### 3.5 Fiber Type Distribution

Results on *FT* are dealt with in their own sub-section because studies in the previous sub-sections did not report *FT* data. Very few studies have reported data from a large number of human lower limb muscles. To the knowledge of the author, only three studies have included data from more than five muscles: Johnson et al. (1973) presented data from 14 lower limb muscles in six cadavers, Garrett et al. (1984) presented data from nine hip and knee muscles in 10 cadavers, and Tirrell et al. (2012) presented data from 36 lower limb muscles in six cadavers. Five additional cadaver studies on fewer muscles were identified: Edgerton et al. (1975), Elder et al. (1982), Parkkola et al. (1993), Dahmane et al. (2005), and Arbanas et al. (2009). A large number of muscle biopsy studies have reported *FT* data from vastus lateralis only; for brevity these studies were excluded from the present summary. Five additional biopsy studies included in the review were: Clarkson et al. (1980), Vandenborne et al. (1993), Plomgaard et al. (2005), Trappe et al. (2009), and Dickinson et al. (2010). Dahmane et al. (2005) reported *FT* as the fraction of fiber cross-sectional area, and Clarkson et al. (1980) and Parkkokla et al. (1993) reported *FT* as both the fraction of fiber count and fiber cross-sectional area; only the fiber count fractions are reported here. All other studies reported *FT* as the fraction of fiber count. Data from Clarkson et al. (1980) are separated into “power” athletes (weightlifters) and “endurance” athletes (runners). In all the data presented here, *FT* was defined as the fraction of muscle fibers that were not counted as “Slow”, “Type I”, or “Myosin Heavy Chain I”.

*FT* data overall are rather sparse in the literature: in the set of 12 studies included here, there were only three muscles with data in more than three studies: gastrocnemius (*N* = 9, mean *FT* = 43.7±9.3%), soleus (8, 22.1±8.5%), and vastus lateralis (8, 55.2±11.7%).

#### 3.6 Muscle Parameters

The full set of muscle parameters from the studies above are included as a Microsoft Excel spreadsheet in the electronic supplementary material. When possible, the data were separated by sex, age, and any other relevant factors in the study such as training status. The coefficient of variation between studies, averaged over muscles and weighted by the number of samples, was 19% for *L_o_*, 62% for *θ*, and 52% for PCSA. Sample results for the quadriceps muscles are presented in Fig. 2: coefficients of variation ranged from 8-11% for *L_o_*, 48-78% for *θ*, and 52-63% for PCSA. Overall coefficients of variation for *L_u_* and *FT* were not calculated due to the relatively small sample sizes of those data.

**Figure 2.**
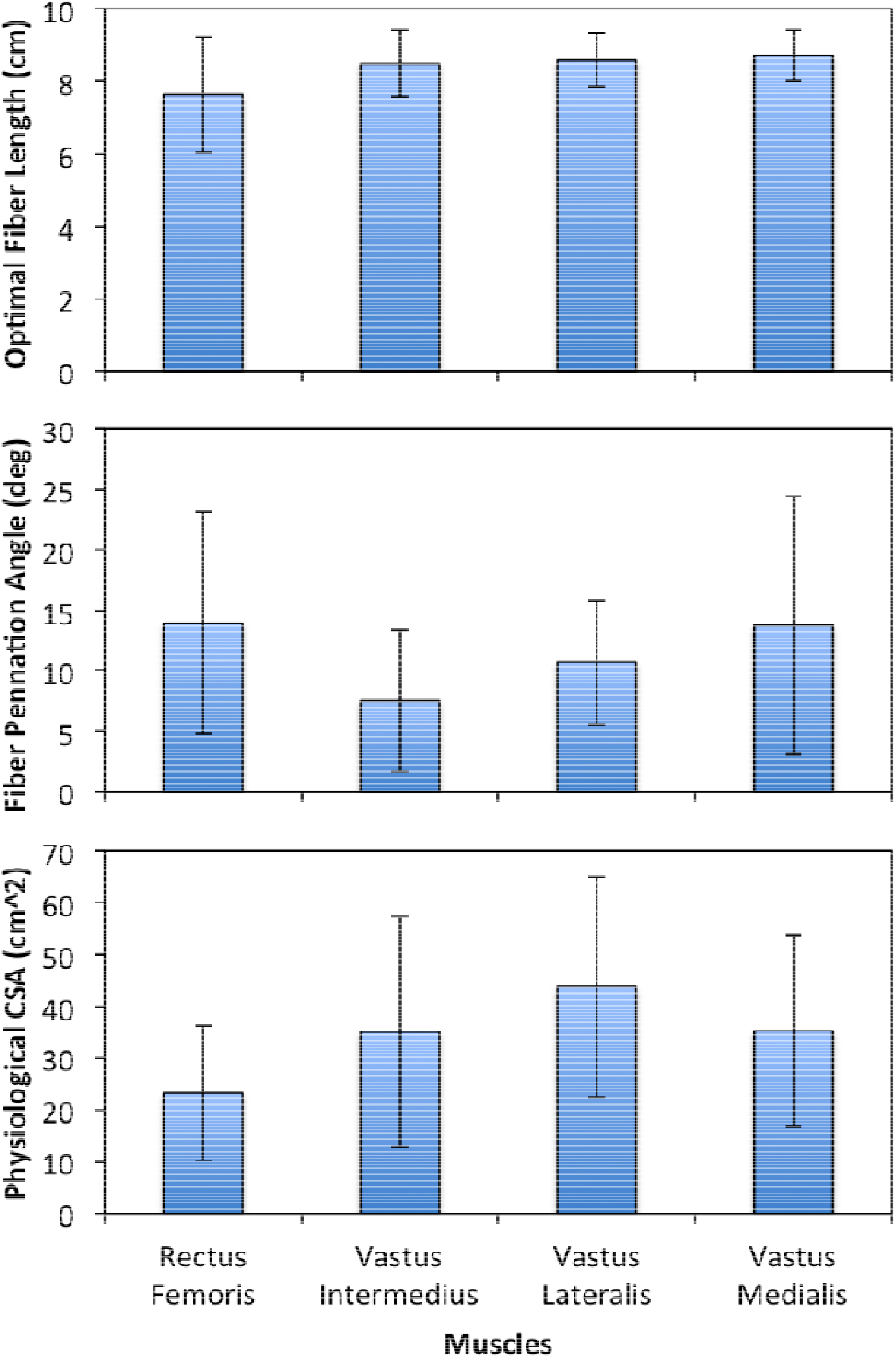
Means and standard deviations (between studies) for optimal fiber lengths, fiber pennation angles, and physiological cross-sectional areas of the quadriceps muscles.

### 4. Discussion

This work summarizes values for five key parameters on Hill-based muscle modeling: the optimal fiber length *L_o_*, the fiber pennation angle *θ*, the PCSA, the unloaded tendon length *L_u_*, and the fast-twitch fiber fraction *FT,* providing a current update to earlier similar efforts on the breadth and depth of these parameter values in the literature (Yamaguchi et al., 1990; van der Helm & Yamaguchi, 2000). Where possible, an effort was made to use consistent practices of fiber length normalization and the definition of PCSA for the data presented here. The review of *FT* includes substantially more data than the earlier summaries. The present results suggest that *L_o_* is remarkably consistent between subjects, but that the other parameters are quite variable. Relatively few studies have reported values for *L_u_* or for any measurement of tendon length, but since whole-muscle length varies widely between subjects, *L_u_* likely also varies widely between subjects especially if the absolute length of *L_o_* is relatively consistent. Given that *L_u_* is arguably the parameter to which Hill model output is most sensitive (e.g. Caldwell, 1995; Scovil & Ronsky, 2006), in situations where subject-specific muscle model parameters may be necessary (e.g. when comparing a patient population to controls), modelers should take care in assigning the value of this parameter and other parameters with high between-subjects variability.

It is hoped that the present summary will be a useful resource for defining muscle model parameters in computer simulations of human movement. A limitation is that the present work does not provide a complete set of reference parameters for Hill-based muscle modeling and musculoskeletal modeling. For example, parameters for the width of the CC force-length relationship were not included, nor were parameters for the CC force-velocity relationship or the SEC force-extension relationship. These parameters are more difficult to summarize than the present parameters because their definitions and methods of implementation in Hill models are substantially more variable. In addition, data for defining muscle paths as functions of skeletal joint kinematics were not included. Some of the cited references here included such data (e.g. Klein Horsman et al., 2007). The availability of software like OpenSim (Delp et al., 2007), where three-dimensional muscle paths can be geometrically scaled and further customized in a user-friendly fashion, has made this important step in musculoskeletal modeling substantially easier.

The need to define muscle paths relates to the use of the parameter values presented here in practice. In a real muscle, “muscle length” *L_M_* refers to the origin-to-insertion length. The analogous length in the Hill model is the CC length plus the SEC length (Fig. 1):

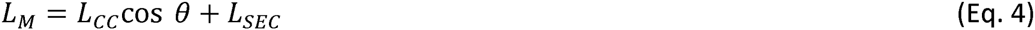

The excursions of *L_M_* will depend on the muscle path definitions and the skeletal kinematics, while the (feasible) excursions of *L_cc_* and *L_SEC_* will depend on the definitions of the force-length and force-extension relationships and the values of *L_o_* and *L_u_* respectively. Directly inserting referenced parameter values for *L_o_* and *L_u_* into a specific musculoskeletal model that exists outside the source of the parameters, with arbitrary muscle path definitions; the musculoskeletal model itself will in nearly all cases need to be directly included in the process of assigning parameter values or vice-versa (e.g. Arnold et al., 2010). As noted earlier, an ideal Hill model will produce the same forces for the same excitation and kinematic conditions as a real muscle; it will not necessarily produce CC/SEC kinematics that match *in* v/Vofiber/tendon kinematics. However, measurements from real muscle should likely form the basis for what reasonable values of Hill model parameters should resemble. For example, although the Hill model is phenomenological in nature, a well-parameterized Hill model of gastrocnemius or soleus should likely not have a relatively long CC / short SEC if it is to be used in simulations of stretch-shortening cycles and the associated work and energy of active muscle contraction (e.g. Lichtwark & Wilson, 2007), even if allowing such parameters gives a better fit with torque-angle dynamometry data than parameters representing a relatively short CC / long SEC.

Hasson and Caldwell (2012) is a rare example where optimization has been used to define muscle model parameters in a specific musculoskeletal model on the basis of matching dynamometry data, using referenced literature values as a starting point. Hatze (1981) was the first to propose this approach in an application in the elbow flexors, and Gerritsen et al. (1998) extended it to the lower limb joints. Other authors have used conceptually similar approaches without formal optimization by hand-tuning selected model parameters to match the general shape of dynamometry data (e.g. Anderson & Pandy, 1999; Arnold et al., 2010). This approach provides useful and reliable estimates of muscle parameters for simulations of whole-body movement (De Groote et al., 2010) and should be strongly considered in any study involving muscle models. In cases where experimental dynamometry data are unavailable or subject-specific parameters are unnecessary, normative data on multiple populations have been published (e.g. Anderson et al., 2007).

A common use of the PCSA data presented here is to calculate the CC maximum isometric force F_0_:

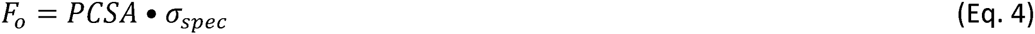

where *σ_spec_* is the specific tension of muscle fibers. The value of *σ_spec_* is in theory constant both within and between subjects, but neither claim has compelling empirical support unless the related literature is reviewed very selectively. Poor agreement on *σ_spec_* between studies could be affected by experimental design, e.g. voluntary vs. electrically-stimulated muscle activation to determine *F_0_* and by differences in determining PCSA (e.g. if intramuscular fat and water content are accounted for). Poor agreement on *o_spec_* between muscles suggests Eq. 4 may by overly simplistic. For example, Eq. 4 does not account for lateral force transmission between fibers (Block & Gonzalez Serratos, 2003), and it assumes pennation angle *θ* is a spatial constant, which is not the case in real muscle (Scott et al., 1993). Human joint torque-angle curves are considerably wider than fiber or sarcomere force-length curves (van den Bogert et al., 1998), suggesting these omissions play an important role in the physiology of muscle force production, and that functional CC lengths are longer than anatomical fiber lengths. This phenomenon further argues for defining parameters like *F_a_* and *L_o_* from joint-level strength tests (e.g. Hasson & Caldwell, 2012) with measurements of fiber/tendon characteristics providing a starting point or the basis for feasible ranges rather than direct parameter values.

## Electronic Supplementary Material

A Microsoft Excel spreadsheet containing the muscle model parameter values is available on Figshare: https://figshare.eom/s/313c3017c6bc2674elc7

## References

Akima H, Kubo K, Imai M, Kanehisa H, Suzuki Y, Gunji A, Fukunaga T (2001). Inactivity and muscle: effect of resistance training during bed rest on muscle size in the lower limb. Acta Physiologica Scandinavica 172, 269–278.

Alexander RM, Vernon A (1975). The dimensions of knee and ankle muscles and the forces they exert. Journal of Human Movement Studies 1,115–123.

Anderson DE, Madigan ML, Nussbaum MA (2007). Maximum voluntary joint torque as a function of joint angle and angular velocity: model development and application to the lower limb. Journal of Biomechanics 40, 3105–3113.

Anderson FC, Pandy MG (1999). A dynamic optimization solution for vertical jumping in three dimensions. Computer Methods in Biomechanics & Biomedical Engineering 2, 201–231.

Arbanas J, Starcevic Klasan G, Nikolic M, Jerkovic R, Miljanovic I (2009). Fibre type composition of the human psoas major muscle with regard to the level of its origin. Journal of Anatomy 215, 636–641.

Arnold EM, Ward SR, Lieber RL, Delp SL (2010). A model of the lower limb for analysis of human movement. Annals of Biomedical Engineering 38, 269–279.

Block RJ, Gonzalez-Serratos H (2003). Lateral force transmission across costameres in skeletal muscle. Exercise & Sport Sciences Reviews 31, 73–78.

Caldwell GE (1995). Tendon elasticity and relative length: effects on the Hill two-component muscle model. Journal of Applied Biomechanics 11,1-24.

Caldwell GE, Chapman AE (1989). Applied muscle modelling: implementation of muscle-specific models. Computers in Biology & Medicine 19, 417–434.

Carbone V, van der Krogt MM, Koopman HFJM, Verdonschot N (2016). Sensitivity of subject-specific models to Hill muscle-tendon model parameters in simulations of gait. Journal of Biomechanics 49,1953–1960.

Clarkson PM, Kroll W, McBride TC (1980). Maximal isometric strength and fiber type composition in power and endurance athletes. European Journal of Applied Physiology 44, 35–42.

Dahmane R, Djordjevič S, Šimunič B, Valenčič V (2005). Spatial fiber type distribution in normal human muscle: histochemical and tensiomyographical evaluation. Journal of Biomechanics 38, 2451–2459.

De Groote F, Van Campen A, Jonkers I, De Schutter J (2010). Sensitivity of dynamic simulations of gait and dynamometer experiments to Hill muscle model parameters of knee flexors and extensors. Journal of Biomechanics 43, 1876–1883.

Delp SL, Anderson FC, Arnold AS, Loan P, Habib A, John CT, Guendelman E, Thelen DG (2007). OpenSim: open-source software to create and analyze dynamic simulations of movement. IEEE Transactions on Biomedical Engineering 54, 1940–1950.

Delp SL, Loan JP, Hoy MG, Zajac FE, Topp EL, Rosen JM (1990). An interactive graphics-based model of the lower extremity to study orthopaedic surgical procedures. IEEE Transactions on Biomedical Engineering 37, 757–767.

Dickinson JM, Lee JD, Sullivan BE, Harber MP, Trappe SW, Trappe TA (2010). A new method to study in vivo protein synthesis in slow-and fast-twitch muscle fibers and initial measurements in humans. Journal of Applied Physiology 108, 1410–1416.

Domire ZJ, Challis JH (2010). A critical examination of the maximum shortening velocity used in simulation models of human movement. Computer Methods in Biomechanics & Biomedical Engineering 13, 693–699.

Elder GCB, Bradbury K, Roberts R (1982). Variability of fiber type distributions within human muscles. Journal of Applied Physiology 53, 1473–1480.

Friederich JA, Brand RA (1990). Muscle fiber architecture in the human lower limb. Journal of Biomechanics 23, 91–95.

Fukunaga T, Roy RR, Shellock FG, Hodgson JA, Day MK, Lee PL, Kwong-Fu H, Edgerton VR (1992). Physiological cross-sectional area of human leg muscles based on magnetic resonance imaging. Journal of Orthopaedic Research 10, 926–934.

Garrett WE, Califf JC, Bassett FH (1984). Histochemical correlates of hamstring injury. American Journal of Sports Medicine 12, 98–103.

Gerritsen KGM, van den Bogert AJ, Hulliger M, Zernicke RF (1998). Intrinsic muscle properties facilitate locomotor control: a computer simulation study. Motor Control 2, 206–220.

Hasson Cj, Caldwell GE (2012). Effects of age on mechanical properties of dorsiflexor and plantarflexor muscles. Annals of Biomedical Engineering 40, 1088–1101.

Hatze H (1976). The complete optimization of a human motion. Mathematical Biosciences 28, 99–135.

Hatze H (1981). Estimation of myodynamic parameter values from observations on isometrically contracting muscle groups. European Journal of Applied Physiology 46, 325–338.

Hatze H (1983). Computerized optimization of sports motions: an overview of possibilities, methods and recent developments. Journal of Sports Sciences 1, 3–12.

Hill AV (1938). The heat of shortening and the dynamic constants of muscle. Proceedings of the Royal Society of London B: Biological Sciences 126, 136–195.

Hill AV (1950). The series elastic component of muscle. Proceedings of the Royal Society of London B: Biological Sciences 137, 273–280.

Johnson MA, Polgar J, Weightman D, Appleton D (1973). Data on the distribution of fiber types in thirty-six human muscles. Journal of Neurological Science 18, 111–129.

Klein Horsman MD, Koopman HFJM, van der Helm FCT, Poliacu Prose L, Veeger HEJ (2007). Morphological muscle and joint parameters for musculoskeletal modelling of the lower extremity. Clinical Biomechanics 22, 239–247.

Koelewijn AD, van den Bogert AJ (2016). Joint contact forces can be reduced by improving joint moment symmetry in below-knee amputee gait simulations. Gait & Posture 49, 219–225.

Lichtwark GA, Wilson AM (2007). Is Achilles tendon compliance optimized for maximum muscle efficiency during locomotion? Journal of Biomechanics 40, 1768–1775.

Lieber RL, Loren GJ, Fridén J (1994). In vivo measurement of human wrist extensor muscle sarcomere length changes. Journal of Neurophysiology 71, 874–881.

Maganaris CN, Baltzopoulos V, Ball D, Sargeant AJ (2001). In vivo specific tension of human skeletal muscle. Journal of Applied Physiology 90, 865–872.

Mendez J, Keys A (1960). Density and composition of mammalian muscle. Metabolism, Clinical & Experimental 9, 184–188.

Narici MV, Maganaris CN, Reeves ND, Capodaglio P (2003). Effect of aging on human muscle architecture. Journal of Applied Physiology 95, 2229–2234.

Navacchia A, Myers CA, Rullkoetter PJ, Shelburne KB (2016). Prediction of in vivo knee loads using a global probabilistic analysis. Journal of Biomechanical Engineering 138, 4032379.

O'Brien TD, Reeves ND, Baltzopoulos V, Jones DA, Maganaris CN (2010). In vivo measurements of muscle specific tension in adults and children. Experimental Physiology 95, 202–210.

Pandy MG (1990). An analytical framework for quantifying muscular action during human movement. In: Winters JM, Woo SLY (eds.), Multiple Muscle Systems, 653–662. Berlin: Springer.

Parkkola R, Alanen A, Kalimo H, Lillsunde I, Komu M, Kormano M (1993). MR relaxation times and fiber type predominance of the psoas and multifidus muscle: an autopsy study. Acta Radiologica 34, 16–19.

Plomgaard P, Penkowa M, Pedersen BK (2005). Fiber type specific expression of TNF-alpha, IL-6 and IL-18 in human skeletal muscles. Exercise Immunology Review 11, 53–63.

Ramsey RW, Street SF (1940). The isometric length-tension diagram of isolated skeletal muscle fibres of the frog. Journal of Cellular & Comparative Physiology 15, 11–34.

Redl C, Gfoehler M, Pandy MG (2007). Sensitivity of muscle force estimates to variations in muscle-tendon properties. Human Movement Science 26, 306–319.

Reeves ND, Narici MV, Maganaris CN (2004). Effect of resistance training on skeletal muscle-specific force in elderly humans. Journal of Applied Physiology 96, 885–892.

Regev GJ, Kim CW, Tomiya A, Lee YP, Ghofrani H, Garfin SR, Lieber RL, Ward SR (2011). Psoas muscle architectural design, in vivo sarcomere length range, and passive tensile properties support its role as a lumbar spine stabilizer. Spine 36, E1666–E1674.

Sacks RS, Roy RR (1982). Architecture of the hindlimb muscles of the cat: functional significance. Journal of Morphology 173, 185–195.

Sanchez GN, Sinha S, Liske H, Chen X, Nguyen V, Delp SL, Schnitzer MJ (2015). In vivo imaging of human sarcomere twitch dynamics in individual motor units. Neuron 88, 1109–1120.

Scott SH, Engstrom CM, Loeb GE (1993). Morphometry of human thigh muscles: determination of fascicle architecture by magnetic resonance imaging. Journal of Anatomy 182, 249–257.

Scott W, Stevens J, Binder-Maclead SA (2001). Human skeletal muscle fiber type classifications. Physical Therapy 81, 1810–1816.

Scovil CY, Ronsky JL (2006). Sensitivity of a Hill-based muscle model to perturbations in model parameters. Journal of Biomechanics 39, 2055–2063.

Thelen DG, Anderson FC (2006). Using computed muscle control to generate forward dynamics simulations of human walking from experimental data. Journal of Biomechanics 39,1107–1115.

Trappe S, Costill D, Gallagher P, Creer A, Peters JR, Evans H, Riley DA, Fitts RH (2009). Exercise in space: human skeletal muscle after 6 months aboard the International Space Station. Journal of Applied Physiology 106, 1159–1168.

Trotter JA (1990). Interfiber tension transmission in series-fibered muscles of the cat hindlimb. Journal of Morphology 206, 351–361.

Tirrell TF, Cook MS, Carr JA, Ward SR, Lieber RL (2012). Human skeletal muscle biochemical diversity. Journal of Experimental Biology 215, 2551–2559.

Umberger BR, Gerritsen KGM, Martin PE (2003). A model of human muscle energy expenditure. Computer Methods in Biomechanics & Biomedical Engineering 6, 99–111.

Van den Bogert AJ, Gerritsen KGM, Cole GK (1998). Human muscle modelling from a user’s perspective. Journal of Electromyography & Kinesiology 8, 119–124.

Van der Helm FCT, Yamaguchi GT (2000). Morphological data for the development of musculoskeletal models: an update. In: Winders JM (ed.), Biomechanics & Neural Control of Posture & Movement, 645–658. Berlin: Springer.

Vandenborne K, Walter G, Ploutz-Snyder L, Staron R, Fry A, De Meirleir K, Dudley A, Leigh JS (1995). Energy-rich phosphates in slow and fast human skeletal muscle. American Journal of Physiology 268, C869–C876.

Walker SM, Schroedt GR (1974). I segment lengths and thin filament periods in skeletal muscle fibers of the Rhesus monkey and the human. Anatomical Record 178, 63–81.

Ward SR, Eng CM, Smallwood LH, Lieber RL (2009). Are current measurements of lower extremity muscle architecture accurate? Clinical Orthopaedics & Related Research 467, 1074–1082.

Weber E (1846). Wagner’s Handworterbuch der Physiologie. Braunschweig: Vieweg.

Wickiewicz TL, Roy RR, Powell PL, Edgerton VR (1983). Muscle architecture of the human lower limb. Clinical Orthopaedics & Related Research 179, 275–283.

Woodley SJ, Mercer SR (2005). Hamstring muscles: architecture and innervation. Cells Tissues Organs 179, 125–141.

Yamaguchi GT, Sawa AGU, Moran DW, Fessler MJ, Winters JM (1990). A survey of human musculotendon actuator parameters. In: Winters JM, Woo SL (eds.), Multiple Muscle Systems, 717–773. Berlin: Springer.

